# *SF3B1*-mutant mis-splicing of *UBA1* confers a targetable therapeutic vulnerability through UBA1 inhibition

**DOI:** 10.1101/2024.08.28.610114

**Authors:** Jonas Thier, Sophia Hofmann, Katharina M Kirchhof, Gabriele Todisco, Teresa Mortera-Blanco, Indira Barbosa, Ann-Charlotte Björklund, André G Deslauriers, Elli Papaemmanuil, Eirini P Papapetrou, Eva Hellström-Lindberg, Pedro L Moura, Vanessa Lundin

**Author notes:** Corresponding Author: Vanessa Lundin, PhD; Address: Department of Medicine, Huddinge, Karolinska Institutet, 141 83 Huddinge, Sweden.

## Abstract

*SF3B1* mutation-driven myelodysplastic syndromes (MDS-*SF3B1*) arise due to somatic mutation in the splicing factor *SF3B1* gene. *SF3B1* mutations induce RNA mis-splicing and loss of expression of critical genes for erythropoiesis, leading to erythroid dysplasia and ultimately refractory anemia. The development of precision medicine approaches for MDS- *SF3B1* is hampered by the complexity of the mis-splicing landscape and its evaluation in disease-accurate model systems. To identify novel RNA mis-splicing events, isogenic *SF3B1*^K700E^ and *SF3B1*^WT^ iPSC lines from an MDS-*SF3B1* patient were differentiated into hematopoietic cells *in vitro* and subjected to unsupervised splicing event analysis using full-length RNA sequencing data. This revealed *SF3B1*^K700E^-specific mis-splicing of ubiquitin-like modifier activating enzyme 1 (*UBA1*) transcripts, which encode the essential E1 protein at the apex of the ubiquitination cascade. *UBA1* mis-splicing (*UBA1*^ms^) preserved *UBA1*^ms^ mRNA but not protein expression. Consequently, *UBA1*^ms^ diminished the pool of functional UBA1, sensitizing *SF3B1*^K700E^ cell lines to the small-molecule UBA1 inhibitor TAK-243. Finally, analysis of CD34^+^ RNA sequencing data from an MDS patient cohort confirmed unique and ubiquitous *UBA1*^ms^ in MDS-*SF3B1* patients, without detection in other splicing factor-mutated MDS patients, or in healthy individuals. TAK-243 selectively targeted MDS-*SF3B1* primary CD34^+^ cells and reduced mutant cell number in colony-forming unit studies. In contrast, normal hematopoietic progenitor cells were unaffected. Altogether, we here define *UBA1*^ms^ as a novel therapeutic vulnerability in *SF3B1*-mutant cells, introducing UBA1 inhibition as a potential avenue for future MDS-*SF3B1* treatments.

## Introduction

Myelodysplastic neoplasms (MDS) are clonal myeloid malignancies that originate in hematopoietic stem cells, characterized by clonal expansion and ineffective hematopoiesis.^1,2^ Heterozygous, recurrent point mutations in splicing factor genes, such as *SF3B1*, *SRSF2*, and *U2AF1* are typically mutually exclusive and the most common variants in MDS patients, resulting in widespread splicing defects.^3,4^

MDS with *SF3B1* mutation (MDS-*SF3B1*) constitute a distinct subgroup of MDS, characterized by bone marrow accumulation of mutant dysplastic erythroblasts, called ring sideroblasts, erythroid cytopenia, and refractory anemia.^5–8^ *SF3B1* encodes subunit 1 of the core RNA splicing factor 3b, involved in processing precursor mRNA to mature transcripts.^9–12^ Missense mutations in this gene alter the interaction of SF3B1 with the pre-mRNA sequence, resulting in extensive cryptic 3’ splice site usage and mis-splicing of genes important for hematopoietic and erythroid differentiation, including *ABCB7*, *ALAS2*, *MAP3K7, PPOX, and TMEM14C*.^13–19^

Therapeutic target studies using experimental disease models are complicated by the dynamic nature of RNA mis-splicing in MDS: mouse models of *SF3B1* mutation develop anemia but feature distinct RNA splicing from humans^20–22^; primary patient material is heterogenous, has limited availability and *in vitro* longevity; and cell models do not fully recapitulate the complex splicing repertoire. Patient-derived induced pluripotent stem cells (iPSCs) are scalable, tractable and genetically faithful models, which are being increasingly utilized to understand myeloid pathobiology.^23–30^

In a previous study, we established isogenic iPSC lines from MDS-*SF3B1* patients and identified a signature of splicing events associated with mutant *SF3B1* in hematopoietic stem and progenitor cells (HSPCs).^23^ Other work has further confirmed a splicing program paralleling the primary disease, supporting the use of *SF3B1*- mutant iPSCs as a physiologically relevant model.^24^

Here, we identify a previously unreported mis-splicing event affecting the gene ubiquitin-activating enzyme 1 (*UBA1*) in the *SF3B1*-mutant context. *UBA1* mis-splicing (*UBA1*^ms^) abrogates protein production, leading to an overall reduction of total UBA1 protein content in *SF3B1*^K700E^ cell lines and an increased sensitivity to the specific UBA1 inhibitor TAK-243. We verified that *UBA1*^ms^ was unique to *SF3B1*-mutant patients in a large MDS cohort,^31^ and similarly conferred a vulnerability to TAK-243 in colony-forming unit assays using primary patient cells. Taken together, we here define *UBA1*^ms^ as a targetable therapeutic vulnerability of mutant cells in MDS-*SF3B1*.

## Results

### *SF3B1* mutation induces *UBA1*^ms^ in MDS-*SF3B1*

New treatment strategies for MDS-*SF3B1* require a deepened understanding of mis-splicing and the molecular consequences in sculpting the pathophysiology of MDS. Here, we employed isogenic iPSC lines originally generated from an MDS*-SF3B1* patient,^23^ harboring an isolated *SF3B1*^K700E^ mutation (**Figure 1A**), the most recurrent variant among MDS*-SF3B1* cases.^10^ To model MDS-*SF3B1 in vitro*, we performed hematopoietic differentiation of wildtype (*SF3B1*^WT^) and mutant (*SF3B1*^K700E^) iPSCs for twelve days and analyzed surface expression of HSPC markers CD34 and CD45 by flow cytometry (**Figure 1B**, left). Consistent with our previous observations, *SF3B1*^K700E^ HSPCs showed lower viability (supplemental **Figure 1A**) but no defects in hematopoietic specification, with similar levels of CD34^+^ cells compared to *SF3B1*^WT^ (supplemental **Figure 1B**). As before, *SF3B1*^K700E^ iPSCs exhibited reduced erythroid potential compared with *SF3B1*^WT^ cells (**Figure 1B**, right).^23^ To examine *SF3B1* mutation-mediated mis-splicing and uncover novel RNA mis-splicing events, we performed 5’-based full-length RNA sequencing of iPSC-derived erythroid cells, to capture both immature and mature, messenger RNA (mRNA) molecules. Splicing analysis validated aberrant splicing of known genes, such as *TMEM14C* and *MAP3K7*, in *SF3B1*^K700E^ but not in *SF3B1*^WT^ cells (supplemental **Figure 1C**), in line with prior observations and *SF3B1*-mutated patients.^16,24,33^ Moreover, in this analysis we also identified an RNA mis-splicing event of the *UBA1* transcript (59.3 percent spliced-in, PSI) caused by an in-frame retention of an intronic region of 135 bases between exon 5 and 6 (**Figure 1C** and supplemental **Figure 1D).** The *UBA1* gene is located on the X-chromosome and expressed in two main isoforms: nuclear UBA1a and cytosolic UBA1b. *UBA1* encodes a multidomain protein that initiates the ubiquitination pathway, which targets proteins for degradation by the ubiquitin-proteasome system through activation and transfer of ubiquitin to E2 enzymes. As *SF3B1* mutations originate in the stem cell population^34^, we performed RT-qPCR for mis-spliced *UBA1* transcripts (supplemental **Figure 1E**) in iPSC-derived HSPCs and erythroid progenitors. *SF3B1*^K700E^ but not *SF3B1*^WT^ cells expressed detectable levels of *UBA1*^ms^ transcript in both hematopoietic cell types (**Figure 1D**). Based on these results, we continued to focus on the HSPC population and investigated whether *UBA1*^ms^ affects protein level in iPSC-derived CD34^+^ by intracellular flow cytometry. Both the percentage of UBA1a positive cells and the calculated staining index were lower in *SF3B1*^K700E^ HSPCs compared with *SF3B1*^WT^ (**Figure 1E**). These results uncover mis-splicing of *UBA1* in an *in vitro* model of MDS-*SF3B1* and suggest a decrease in UBA1 protein content.

**Figure 1.**
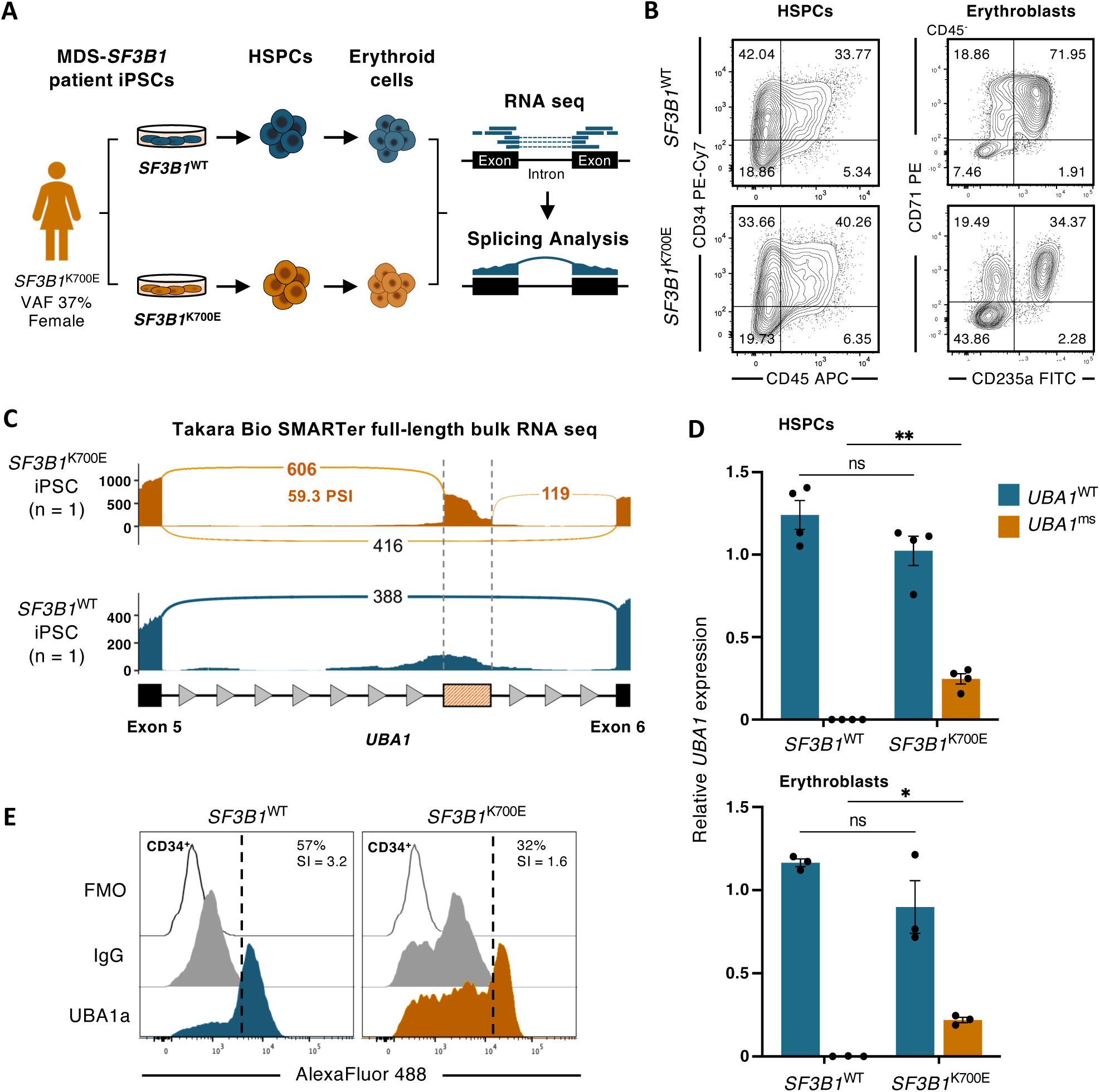
RNA sequencing reveals *UBA1*^ms^ in an iPSC model of MDS-*SF3B1*. (A) Origin of iPSC lines and experimental overview. (B) Representative flow cytometry diagrams of *SF3B1*^WT^ and *SF3B1*^K700E^ iPSC-derived cells after 12 days of hematopoietic differentiation (left) and another 14 days of erythroid culture (right). (C) Sashimi plots of the mis-spliced region of *UBA1* in *SF3B1*^WT^ and *SF3B1*^K700^ from total RNA sequencing of iPSC-derived GlyA^+^ erythroblasts (n = 1). Black, canonical splice junction counts; orange, mis-spliced junction counts. y-axis, absolute read counts. (D) qPCR analysis of canonically and mis-spliced *UBA1* transcripts (*UBA1*^WT^ and *UBA1*^ms^, respectively), normalized to total *UBA1* in HSPCs (n = 4) and erythroblasts (n = 3) derived from *SF3B1*^WT^ and *SF3B1*^K700E^ iPSCs. Mean ± SEM, relative fold change. Unpaired t test with Holm-Šídák’s multiple comparisons test (E) Intracellular flow cytometry diagrams of UBA1a signal in CD34^+^ iPSC-HSPCs from *SF3B1*^WT^ and *SF3B1*^K700E^. Positive events were gated from IgG negative controls and staining indexes (SI) calculated from AlexaFluor 488 MFI of UBA1- and IgG-labelled populations. *, *P* ≤ 0.05; **, *P* ≤ 0.01; ns, not significant. VAF, variant allele frequency; RNA seq, RNA sequencing.

### *UBA1* RNA mis-splicing compromises protein production

RNA mis-splicing by mutant *SF3B1* may lead to nonsense-mediated decay (NMD) of transcripts or production of aberrant proteins.^13^ To determine the consequence of *UBA1*^ms^ on protein synthesis and function, we first modelled its structure *in silico*. AlphaFold2 was used to predict protein folding of the 1058 amino acid UBA1a isoform from full-length *UBA1* (*UBA1*^WT^) cDNA (ENST00000377351.8) and *UBA1*^ms^ including the predicted 45 amino acid sequence (supplemental **Figure 1D**). The insertion resulted in low sequence coverage, low plDTT (< 50%), and high predicted aligned error (supplemental **Figure 2A-C**). Consequently, the model failed to confidently predict folding of amino acid residues from *UBA1*^ms^ (supplemental **Figure 2D**).

To experimentally test whether *UBA1*^ms^ compromises protein production, we designed expression plasmids for *UBA1*^WT^ (WT) and *UBA1*^ms^ (MS) fused to a C-terminal FLAG tag for transfection of K562 cells and subsequent assessment of recombinant protein levels (**Figure 2A** and supplemental **Figure 3A**). Efficient transfection was confirmed at 72 hours by eGFP positivity using flow cytometry (supplemental **Figure 3B**). *UBA1*^ms^ transcription was first measured by RT-qPCR in cells transfected with WT or MS plasmid or empty vector control. As expected, *UBA1*^ms^ transcript was detected in cells overexpressing MS but not WT or control (**Figure 2B**). To examine whether *UBA1*^ms^ is translated into protein, we performed immunoblotting against UBA1a/b and FLAG 72 hours post-transfection. Endogenous UBA1 protein was present in all conditions (**Figure 2C**, top, and **2D**). However, recombinant FLAG-tagged UBA1 was only detected in cells transfected with WT but not MS or control (**Figure 2C**, middle, and **2D**). To elucidate whether the lack of UBA1 protein may be due to rapid turnover, cells were treated with proteasome inhibitor MG-132 or ATF6α unfolded protein response pathway inhibitor Ceapin-A7. Inhibition of these protein degradation pathways did not result in detectable levels of protein from *UBA1*^ms^ by immunoblotting (**Figure 2C-D**). This was additionally confirmed by immunoprecipitation of FLAG-tagged protein and immunoblotting for FLAG and UBA1a/b, in which significant pull-down of recombinant UBA1 was detected in cells transfected with WT but not MS or control (**Figure 2E-F**). Together, these results show that *UBA1*^ms^ generates persistent transcripts but fails to produce protein, suggesting a reduction in the pool of UBA1 protein.

**Figure 2.**
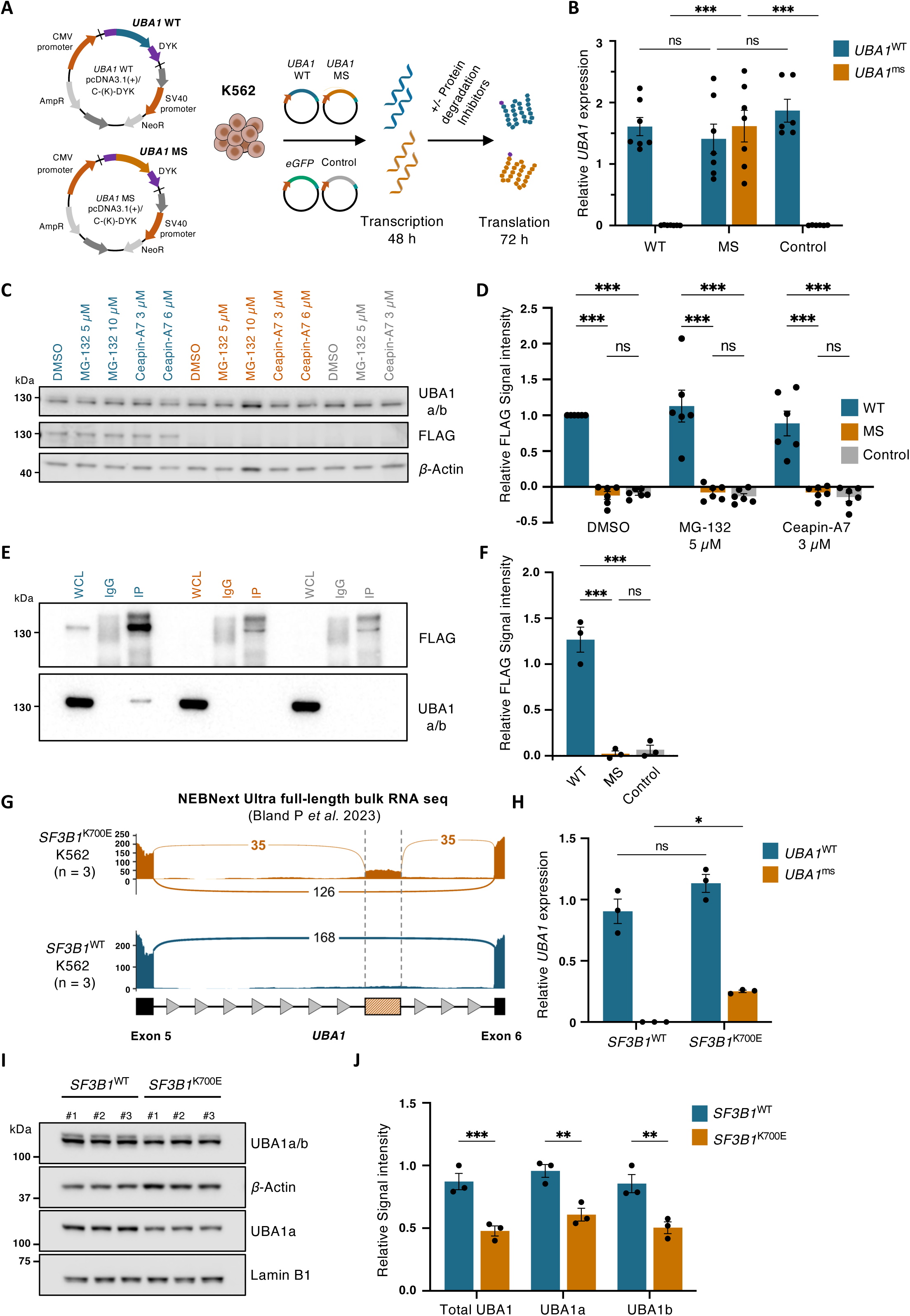
*SF3B1*^K700E^-specific *UBA1*^ms^ compromises protein translation in K562 cells. (A) Maps for ORF cDNA in pcDNA3.1+/C-(K)-DYK plasmids for *UBA1* transcript variant 1 (WT) and *UBA1*^ms^ (MS) and experimental design for evaluation of *UBA1*^ms^ in K562 cells. (B) qPCR analysis of canonically spliced *UBA1*^WT^ and *UBA1*^ms^ transcripts relative to total *UBA1*, in K562 cells 48 hours following transfection with WT (n = 7), MS (n = 7) or pcDNA3.1 control (n = 6). Mean ± SEM, relative fold change. Two-way ANOVA with Tukey’s multiple comparisons test. (C) Representative immunoblot analysis of UBA1a/b and FLAG-tagged protein and (D) quantification of FLAG-tagged protein in K562 cells 72 hours after transfection with WT, MS or control. Cells were treated with MG-132, Ceapin-A7 or DMSO for 24 hours. ß-actin was used as a loading control to normalize results (n = 6). Mean ± SEM, relative FLAG signal intensity. Two-way ANOVA with Tukey’s multiple comparisons test. (E) Representative immunoblot analysis and (F) quantification of immunoprecipitation of FLAG-tagged protein from K562 cells 72 hours after transfection with WT, MS or control. Sample loading scheme: whole cell lysate (WCL), immunoprecipitation by mouse IgG (IgG) and immunoprecipitation by anti-FLAG antibody (IP) (n = 3). Mean ± SEM relative FLAG signal intensity, One-way ANOVA with Tukey’s multiple comparisons test. (G) Sashimi plots of the *UBA1*^ms^ region from total RNA sequencing of *SF3B1*^WT^ and *SF3B1*^K700E^ K562 cells from previously published data.^35^ (H) qPCR analysis of canonically spliced *UBA1* and *UBA1*^ms^ transcripts, relative to total *UBA1*, in *SF3B1*^WT^ and *SF3B1*^K700E^ K562 cells (n = 3). Mean ± SEM relative fold change. Unpaired t test with Holm-Šídák’s multiple comparisons test. (I) Representative immunoblot analysis of UBA1 isoforms and (J) quantification of whole cell lysates from *SF3B1*^WT^ and *SF3B1*^K700E^ K562 cells (n = 3). ß-actin was used to normalize total UBA1 and UBA1b; Lamin B1 was used to normalize UBA1a. Mean ± SEM relative signal intensity. Two-way ANOVA with Šídák’s multiple comparisons test. *, *P* ≤ 0.05; **, *P* ≤ 0.01, ***, *P* ≤ 0.001; ns, not significant.

We next utilized the *SF3B1*^K700E^ K562 human erythroleukemia cell line to examine effects of *UBA1*^ms^ at steady state. Similar to iPSCs, K562 cells harboring *SF3B1* mutation have shown to recapitulate the splicing landscape of primary MDS.^17,19^ First, analysis of a recently published full-length RNA sequencing dataset also displayed the *UBA1*^ms^ event in *SF3B1*^K700E^ but not in *SF3B1*^WT^ K562 cells, which was not reported by the authors (**Figure 2G**).^35^ Again, RT-qPCR analysis of *UBA1* splice forms experimentally confirmed that *UBA1*^ms^ is specific to *SF3B1*^K700E^ K562 cells (**Figure 2H**). To assess the effect of *UBA1*^ms^ on protein level, we performed immunoblotting with antibodies against UBA1a/b and UBA1a isoforms (**Figure 2I**). Total UBA1 protein was significantly decreased in *SF3B1*^K700E^ cells compared to *SF3B1*^WT^, resulting from lower levels of both UBA1a and UBA1b in mutant K562 cells (**Figure 2J**). Notably, we did not detect a higher molecular weight band corresponding to the theoretical 45 amino acids larger protein that would result from translation of *UBA1*^ms^ transcript with either antibody. In summary, these data demonstrate *SF3B1* mutation-specific *UBA1*^ms^ and confirm a decrease in total UBA1 protein levels in mutant cells.

### *SF3B1* mutation sensitizes K562 cells to UBA1 inhibition

UBA1 plays a critical role in cellular proteostasis and inhibition of *UBA1* induces cell death,^36–38^ indicating a minimum level of UBA1 activity is required for cell survival. We hypothesized that a reduction in UBA1 protein content in *SF3B1*-mutant cells would confer higher sensitivity to targeted UBA1 inhibition compared to WT *SF3B1* cells. To test this, the selective UBA1 inhibitor TAK-243^39^ was employed to investigate the effect of UBA1 inhibition on cell viability, clonal competition, and CFU potential in *SF3B1*-mutated cells (**Figure 3A**). We treated K562 cells with increasing doses of TAK-243 and confirmed an inverse decrease in K48-linked ubiquitin, as a proxy for global ubiquitination, (supplemental **Figure 4A)** and determined LD_50_ by flow cytometry after 24 hours (supplemental **Figure 4B**). TAK-243 induced apoptosis in a dose-dependent manner and importantly, with significantly higher sensitivity of *SF3B1*^K700E^ K562 cells compared to WT cells (**Figure 3B-C**). We next tested whether the difference in sensitivity to TAK-243 may be exploited to specifically target *SF3B1*- mutated cells in K562 co-culture at a 50:50 ratio of WT to mutants by measuring VAF by ddPCR after 72 hours of UBA1 inhibition. Indeed, TAK-243 significantly reduced *SF3B1*^K700E^ mutational burden in co-cultures (**Figures 3D**), albeit with modest effect, likely attributed to the proliferative advantage of *SF3B1*^WT^ cells *in vitro*^23^ (supplemental **Figure 4C**), as supported by a reduction in *SF3B1*^K700E^ VAF after 72 hours also in DMSO-treated conditions (**Figure 3D**).

**Figure 3.**
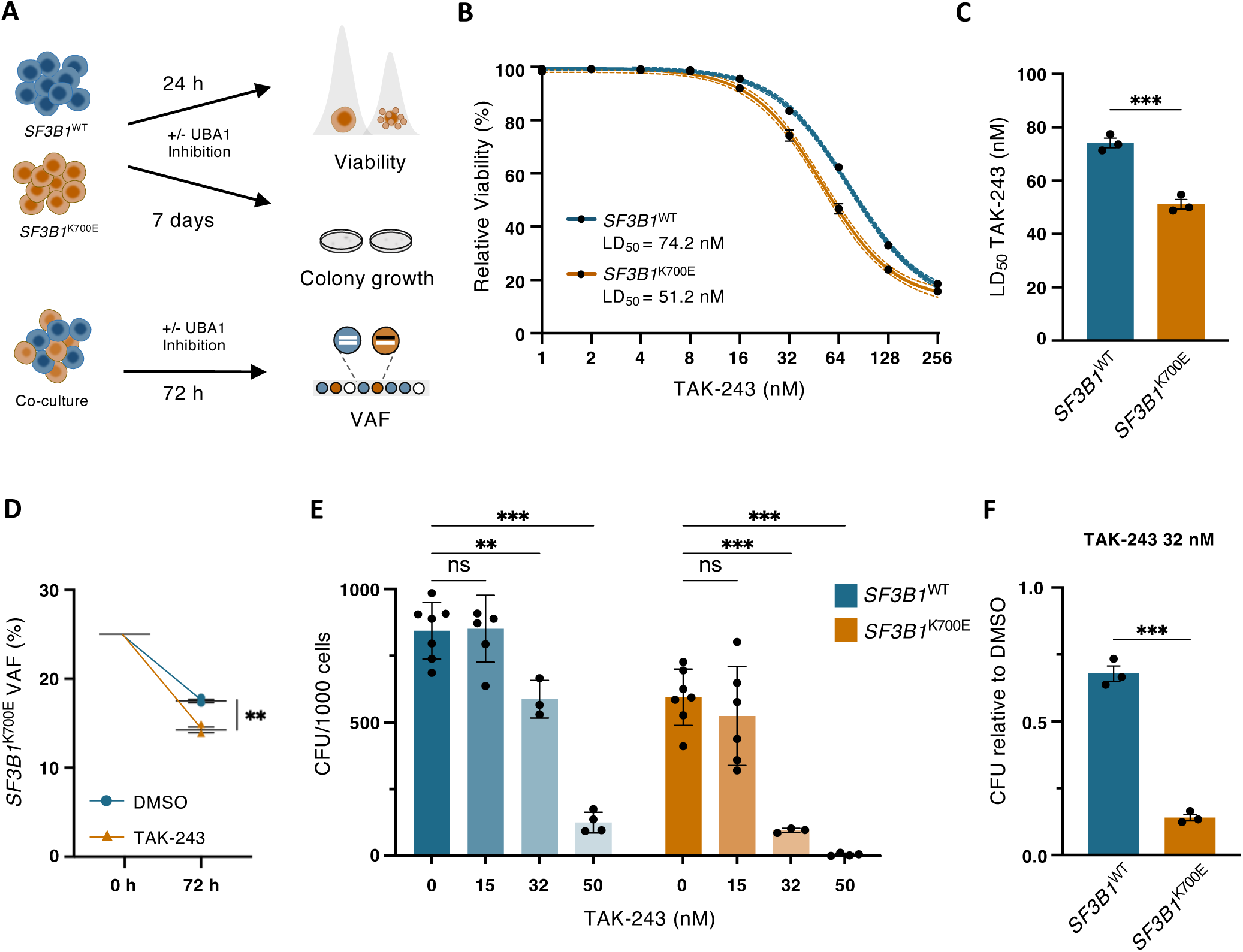
*SF3B1* mutation sensitizes K562 cells to UBA1 inhibition. (A) Experimental strategy to assess the effect of targeted UBA1 inhibition by TAK-243 treatment on *SF3B1*^WT^ and *SF3B1*^K700E^ cells. (B) Dose response curves of *SF3B1*^WT^ and *SF3B1*^K700E^ K562 cells treated with TAK-243 or DMSO for 24 hours (n = 3). Data points represent mean ± SEM live cell fractions, normalized to control-treated samples. Interpolated sigmoidal, 4PL, standard curves of mean (solid line) and 95% confidence interval (dotted line). Live cells were defined as Aqua/Apotracker Green double negative singlets, assessed by flow cytometry with two technical repeats per condition. LD_50_ of TAK-243 displayed under the curves are quantified in (C). Mean ± SEM, nM TAK-243. Unpaired t test. (D) *SF3B1*^K700E^ VAF in co-cultures of 1:1 *SF3B1*^WT^ and *SF3B1*^K700E^ K562 cells after 72 hours of treatment with 50 nM TAK-243 or DMSO, as determined by ddPCR (n = 3). Mean ± SEM. Unpaired t test. (E) CFU counts per 1000 seeded cells. Mean ± SEM. Two-way ANOVA with Dunnett’s multiple comparisons test. And (F) CFU counts relative to DMSO-treated controls, of *SF3B1*^WT^ and *SF3B1*^K700E^ K562 cells treated with increasing concentrations of TAK-243 for 7 days (n^DMSO^ = 7, n^15^ ^nM^ = 5, n^32^ ^nM^ = 3; n^50^ ^nM^ = 4). Mean ± SEM. Unpaired t test. **, P ≤ 0.01; ***, P ≤ 0.001; ns, not significant.

To further investigate the distinct sensitivity of *SF3B1*-mutated cells to UBA1 inhibition, we conducted CFU assays of K562 cells in the presence of TAK-243. We found that the overall CFU potential of *SF3B1*^K700E^ cells was lower than *SF3B1*^WT^ (**Figure 3E**), in line with previous results.^23^ Moreover, UBA1 inhibition affected CFU potential of both *SF3B1*^K700E^ and *SF3B1*^WT^ cells in a dose-dependent manner: while 15 nM TAK-243 exposure had a minimal effect on colony number compared to DMSO control, 50 nM TAK-243 eradicated colonies of mutants but also largely of WT cells (**Figure 3E**). However, treatment with 32 nM resulted in a significantly differential effect on CFU potential, with enhanced sensitivity to TAK-243 of *SF3B1*-mutated cells compared to WT (**Figure 3E-F**). Thus, our results demonstrate a therapeutic window in which *SF3B1*^K700E^ but not *SF3B1*^WT^ cells are sensitized to UBA1 inhibition, which can be harnessed to specifically diminish the number of *SF3B1*-mutant cells while sparing WT clones.

### *UBA1*^ms^ in MDS-*SF3B1* patients confers sensitivity to targeted UBA1 inhibition

We further probed the increased sensitivity of *SF3B1*-mutated cells to UBA1 inhibition by treating patient iPSC-HSPCs with increasing doses of TAK-243 over 24 hours and measuring cell viability by flow cytometry (**Figure 4A** and supplemental **Figure 5A**). As before, TAK-243 was more potent at inducing cell death in *SF3B1*^K700E^ iPSC-HSPCs compared to *SF3B1*^WT^ cells (supplemental **Figure 5B**), resulting in a significantly lower LD_50_ in *SF3B1*^K700E^ HSPCs (**Figure 4B**). Importantly, splicing analysis of RNA sequencing data from bone marrow mononuclear cells (BM MNC) of the original MDS-*SF3B1* patient used in the iPSC reprogramming^23^ validated that *UBA1*^ms^ is a shared event by the mutant cell line and the primary cells (**Figure 4C**). To evaluate the clinical relevance of our findings, we performed splicing analysis in our previously studied MDS cohort,^31^ to verify how *UBA1*^ms^ is recapitulated in patients. Specifically, we explored the mis-splicing event in 124 patients with MDS with ring sideroblasts (*SF3B1*^mt^, n = 83; *SRSF2*^mt^, n = 15; *U2AF1*^mt^, n = 4 and splicing factor^WT^, n = 22) compared to healthy donors (n = 16) using full-length total RNA sequencing data from CD34^+^ BM MNC (**Figure 4D**). *UBA1*^ms^ was abundant (57.31 ± 1.78 PSI) and exclusive to *SF3B1*-mutated patients but not present in the other splicing factor-mutated or WT cases nor healthy donors (**Figure 4E**). To validate these results, we assessed an independent long-read single-cell RNA sequencing dataset.^40^ Although the *UBA1*^ms^ event was not originally reported, splicing analysis confirmed it to be *SF3B1* mutation-exclusive (supplemental **Figure 6A**). This long-read transcriptomic data also enabled verification that *UBA1*^ms^ results in a full-length transcript. Total *UBA1* transcript levels were not significantly different between CD34^+^ cells from sex-matched MDS-*SF3B1* cases and healthy donors. However, *UBA1* transcript levels were overall higher in female compared to male individuals (**Figure 4F)**, likely due to an escape from X inactivation.^41^ *SF3B1*^K700E^ patients, which were the most represented (59%) among the *SF3B1* mutation cohort, exhibited similar levels of *UBA1*^ms^ compared to all other *SF3B1* variants (supplemental **Figure 6B**) with equally high levels of mis-splicing across *SF3B1*^K700E^ VAFs (supplemental **Figure 6C**).

**Figure 4.**
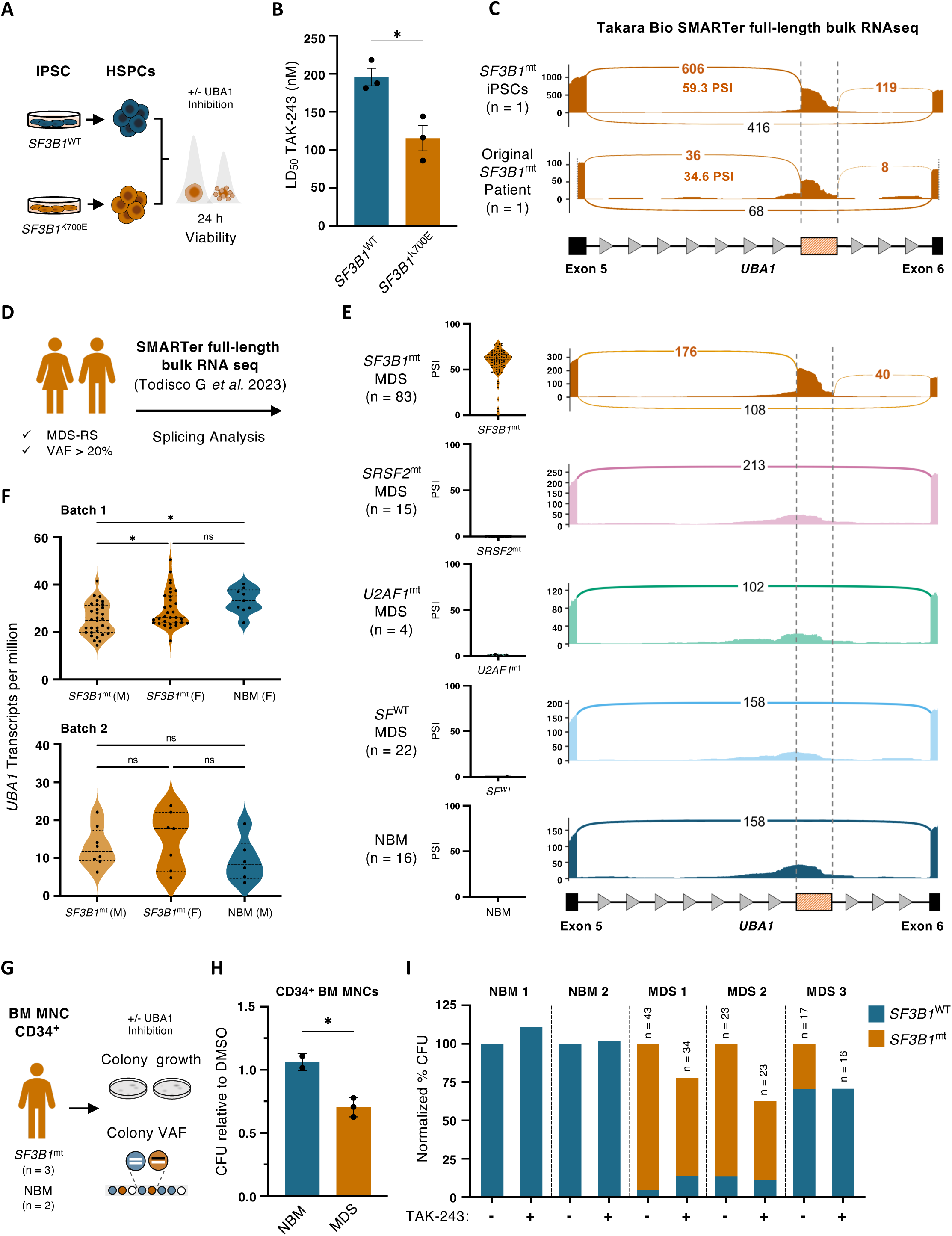
*UBA1*^ms^ in MDS-*SF3B1* patients confers sensitivity to UBA1 inhibition. (A) Experimental strategy to assess the effect of UBA1 inhibition by TAK-243 on the viability of HSPCs derived from *SF3B1*^WT^ and *SF3B1*^K700E^ iPSCs. (B) LD_50_ of TAK-243 in *SF3B1*^WT^ and *SF3B1*^K700E^ iPSC-derived HSPCs treated with TAK-243 or DMSO for 24 hours (n = 3). Mean ± SEM. Unpaired t test. (C) Sashimi plots of the mis-spliced region in *UBA1* from total RNA sequencing of *SF3B1*^K700E^ iPSC-derived erythroblasts and CD34^+^ BM MNCs of the MDS-*SF3B1* patient of iPSC origin. (D) Cohort selection for the splicing analysis of full-length RNA sequencing of CD34^+^ cells from BM MNCs of MDS-RS patients. (E) Violin plots of PSI *UBA1* intron 5 mis-splicing from total RNA sequencing of CD34^+^ BM MNCs, in the cohort of MDS-RS patients and healthy donors, grouped by splicing factor mutation status (left) and sashimi plots of read counts of the mis-spliced region in *UBA1* in patients by splicing factor mutation (n*^SF3B1^*= 83, n*^SRSF2^*= 15, n*^U2AF1^*= 4, n^SFWT^ = 22, n^NBM^ = 16; right). (F) Violin plots of *UBA1* transcript per million counts in *SF3B1*-mutated and NBM samples from the sequenced cohort separated by sequencing batch (Batch 1: *SF3B1*^mt^ (M), n = 32, *SF3B1*^mt^ (F), n = 32, NBM (F), n= 9; Batch 2: *SF3B1*^mt^ (M), n = 8, *SF3B1*^mt^ (F), n = 7, NBM (F), n= 6). One-way ANOVA with Tukey’s multiple comparisons test. (G) Experimental strategy to assess the effect of UBA1 inhibition on colony growth and composition in CD34^+^- enriched BM MNCs from *SF3B1*^mt^ patients and healthy controls. (H) Effect of UBA1 inhibition on CFU counts relative to DMSO and (I) frequency of *SF3B1*^WT^ and *SF3B1*^mt^ colonies relative to total CFU counts from MDS patient (n = 3) or healthy control (n = 2) cells treated with 32 nM TAK-243 or DMSO for 14 days. Numbers indicate colonies assessed by ddPCR. Mean ± SEM. Unpaired t test. *, P ≤ 0.05; ns, not significant. *SF3B1*^mt^*, SF3B1-*mutated; MDS-RS, MDS with ring sideroblasts; NBM, normal bone marrow from healthy donors; M, male; F, female.

Based on the abundance of *UBA1*^ms^ in MDS-*SF3B1* patients in conjunction with the lack of detectable protein from ectopic overexpression of *UBA1*^ms^, we hypothesized a partial loss of functional UBA1 protein that could be exploited therapeutically. To assess whether targeted UBA1 inhibition through TAK-243 in primary HSPCs from MDS patients would selectively reduce *SF3B1*-mutant cells, as observed with K562 cells, we performed CFU assays with CD34^+^ BM MNCs from sex-matched *SF3B1*- mutated MDS patients and healthy donors (**Figure 4G**). Indeed, TAK-243 exposure significantly reduced the CFU potential relative to DMSO of cells from MDS patients compared to healthy donors (**Figure 4H**). Single-colony genotyping by ddPCR confirmed that the treatment effect in *SF3B1*-mutated patient cells largely resulted from a reduction of mutated colonies, whereas CFU formation from residual WT cells was less affected (**Figure 4I**). Taken together, we confirm extensive *UBA1*^ms^ as exclusive to *SF3B1*-mutated patients, conferring a sensitivity of mutant cells to targeted inhibition of UBA1 with TAK-243 in both iPSC-derived and primary HSPCs.

## Discussion

A detailed understanding of mis-splicing and ensuing molecular consequences in splicing factor-mutated MDS is key for the development of improved therapies. Here, we have capitalized on *in vitro* models of MDS-*SF3B1* and identified *UBA1* as a target of mis-splicing. This finding was corroborated by MDS patient cohort analyses, in which *UBA1*^ms^ was exclusively present in *SF3B1-*mutant individuals. Our study supports the relevance of using iPSCs as a tool in hematological research and uncovers an RNA mis-splicing event by mutant *SF3B1* with therapeutic implications. SF3B1 is the most frequently mutated component of the spliceosome machinery, associated with widespread disruption of RNA splicing.^3,8,12,13,16,18,21^ Previously characterized *SF3B1*-mutant mis-spliced transcripts typically introduce a premature stop codon and are therefore targeted for NMD. This was not the case for *UBA1*^ms^, which instead encoded a 45 amino acid insertion within the inactive adenylation domain of UBA1. A recent study suggested *UBA1* RNA and protein expression are decreased in splicing gene mutant MDS, albeit without a link to RNA mis-splicing of *UBA1*.^42^ In contrast, we did not observe differences in *UBA1* transcript expression level between *SF3B1*^mt^ individuals and healthy donor and attribute the reduction of UBA1 protein to the *UBA1*^ms^ event, which did not produce protein in our *in vitro* models. Indeed, *UBA1*^ms^ was detected in existing long-read RNA sequencing data,^40^ demonstrating the generation of mature mRNA from mis-spliced transcripts.

The mechanism by which *UBA1*^ms^ leads to protein deficiency is speculative but our results suggest it is independent of proteasomal degradation. It is possible that the retained intronic sequence induces a conformational change of *UBA1*^ms^ RNA, abrogating translation through ribosome stalling. Characterization of secondary RNA structures due to *UBA1*^ms^ may introduce an additional layer of targeted treatment intervention.

Previous work by us and others has described stage-specific, pervasive mis-splicing due to *SF3B1* mutation and induction of pro-survival mechanisms in mutated cells.^8,40^ In contrast, lower UBA1 protein levels due to *UBA1*^ms^ would likely challenge mutant cell viability as UBA1 is essential for protein homeostasis and cell survival.^36,37^ Curiously, a recent study reported an adaptive stress response in cells with a partial reduction in UBA1 activity that confers cellular resilience^43^, which would be in line with the clonal advantage of *SF3B1*-mutated hematopoietic stem cells.

MDS-*SF3B1* patients frequently develop transfusion-dependency, and allogenic stem cell transplantation remains the only curative option. Advances in single-cell and integrative genomics have empowered the discovery of novel MDS-*SF3B1* disease mechanisms, presenting potential therapeutic modalities.^8,19,40^ Here, we leveraged full-length RNA sequencing data with an unsupervised analysis approach to uncover a new mis-splicing event and demonstrated adverse effects on UBA1 protein levels. Somatic mutation in the *UBA1* gene has recently gained attention in VEXAS (vacuoles, E1 enzyme, X-linked, autoinflammatory, somatic) syndrome, leading to a loss of cytosolic UBA1b and expression of a catalytically impaired isoform.^44^ Symptoms include inflammatory and hematological conditions and interestingly, about half of VEXAS patients also present with MDS.^45^ However, the shared convergence of UBA1 as a target of mis-splicing in MDS-*SF3B1* and somatic mutation in VEXAS, leads to two distinct clonal disorders. Despite the different clinical phenotypes, it is tempting to speculate that exploiting UBA1 dysfunction might be a plausible treatment strategy in both myeloid disorders. Indeed, a recent study of a VEXAS cell model demonstrated selective susceptibility to UBA1 inhibition.^38^ In line with this, our data reveal a differential response of *SF3B1*-mutated cells to TAK-243 compared to WT, giving support to the specificity and applicability of UBA1 inhibition in the development of pharmacological intervention for MDS-*SF3B1*. Utilizing a cell line model as well as HSPCs from iPSCs and MDS patients, we further demonstrate cell type-specific and dose-dependent responses to UBA1 inhibition. Importantly, mutant cells had increased sensitivity to UBA1 inhibition compared with WT cells. Interestingly, TAK-243 has been shown to induce apoptosis in several hematological cancer models, including acute myeloid leukemia,^46–48^ and is currently in clinical trial for intermediate-2 or high-risk refractory MDS and leukemias (NCT03816319). In conclusion, we identify *UBA1* as a previously unrecognized target of *SF3B1*-mutant mis-splicing, which provides specific vulnerability of *SF3B1*-mutated cells to directed UBA1 inhibition.

## Methods

### iPSC culture

The generation and characterization of human iPSC lines MDS-22.44 (*SF3B1*^K700E^) and N-22.45 (*SF3B1*^WT^), from a female MDS patient with ring sideroblasts, single-lineage dysplasia and an isolated *SF3B1*^K700E^ variant, were previously described.^23^ iPSCs were cultured on Matrigel hESC-Qualified Matrix (Corning) in mTeSR Plus (StemCell Technologies) and clump passaged with EZ-LiFT Stem Cell Passaging Reagent (Sigma-Aldrich). HSPCs were generated using STEMdiff Hematopoietic Kit (StemCell Technologies) and harvested on day 12. Methods for erythroid differentiation and RNA sequencing can be found in supplemental Methods.

### Erythroid differentiation and RNA sequencing

Hematopoietic differentiation was adapted from Matsubara *et al*. (PMID: 30948156): iPSCs were seeded as clumps and differentiated in Essential 8 Medium (Gibco), 1% P/S, 80 ng/mL VEGF, 80 ng/mL BMP4, and 2 µM CHIR-99021 from day 0; Essential 6 Medium (Gibco), 1% P/S, 80 ng/mL VEGF, 50 ng/mL SCF, and 2 µM SB431542 from day 2; and Stemline II (Sigma-Aldrich), 1% P/S, 50 ng/mL SCF, 50 ng/mL FLT3L, and 50 ng/mL IL3 from day 4-13 with media changes every other day. For erythroid specification, cells were cultured for 14 days in StemPro-34 SFM with 1% Bovine Albumin Fraction V (Gibco), 1% P/S, 2 mM L-Glutamine, 3.5 µM 1-Thioglycerol, 150 µg/mL holo-Transferrin (Sigma-Aldrich), 2 U/mL Erythropoietin (Retacrit, Pfizer), 50 ng/mL SCF and 50 ng/mL IL3, which was omitted from day 8, with media changes every other day. All cytokines were purchased from PeproTech; and CHIR99021 and SB431542 from StemCell Technologies. Glycophorin A^+^ cells were enriched using CD235a Microbeads and positive selection with autoMACS Pro Separator (Miltenyi Biotec). Automated RNA extraction was performed using QIAcube Connect with RNeasy Micro Kit (Qiagen). Full-length bulk RNA sequencing and analysis was performed as described below.

### RNA splicing analysis

RNA sequencing data was assessed as previously described.^8^ Briefly, differential splicing analysis was performed using rMATS v. 4.1.1, with p-values calculated using the likelihood-ratio test (LRT) and adjusted with the BH method. Sashimi plots for visualization were generated using ggsashimi v. 1.1.5.

### Primer Design

For splice form quantification, primer design was adapted from (PMID: 26979160) to generate primer pairs spanning the normal splice junction (canonical), priming within the mis-spliced sequence (variant) and upstream of the splice site (external control) using Primer-BLAST primer design tool (https://www.ncbi.nlm.nih.gov/tools/primer-blast/). Primers had to meet default requirements with the following modifications: Primers must span an exon-exon junction with a product range between 70-200 nucleotides, a GC content from 40-60%, and include a GC clamp. Primer pair specificity was analyzed using the *Homo sapiens* Refseq mRNA database. Primers had at least four total mismatches, including at least three within the last four 3’ base pairs and a maximum amplicon size.

### Quantitative reverse transcription PCR (RT-qPCR)

Cells were lysed in 350 µL Buffer RLT Plus (Qiagen) containing 40 mM 1,4-Dithiothreitol (DTT; Sigma-Aldrich). Automated RNA extraction was performed using QIAcube Connect with RNeasy Mini Kit (Qiagen) and RNA yield and purity measured by Nanodrop 2000 Spectrophotometer (Thermo Scientific). cDNA was generated using Maxima First Strand cDNA Synthesis Kit for RT-qPCR (Thermo Scientific) from 200 ng RNA. 4 ng cDNA was analyzed in triplicate by two-step reverse transcription quantitative PCR (RT-qPCR) using PowerUp SYBR Green and StepOnePlus Real-Time PCR System (Applied Biosystems). All RT-qPCR experiments included non-template negative controls. *UBA1* splice variant was quantified by fold change expression relative to total *UBA1* external control and *18S* as housekeeping gene. Primer specifics are listed in Supplemental Table 1.

### *UBA1* Expression plasmids

Human *UBA1* transcript variant 1 (WT) cDNA open reading frame (ORF) clone (accession number: NM_003334.4, CloneID: OHu24932) in a pcDNA3.1+/C-(K)-DYK expression vector; pcDNA3.1+C-eGFP (eGFP); and pcDNA3.1(+) (empty vector control) were purchased from GenScript. *UBA1*^ms^ cDNA ORF clone including the 135 extra bases was generated and subcloned into UBA1_OHu24932D_pcDNA3.1+/C-(K)-DYK using GenScript gene synthesis and subcloning service.

### Transfection of K562 cells

K562 cells were maintained in RPMI 1640 Medium with 10% heat inactivated Fetal Bovine Serum (FBS; Gibco) and 1% Penicillin-Streptomycin (P/S, HyClone) with media changes every other day. Cells were passaged two days before transfection: 2×10^5^ cells were nucleofected with 1 µg of *UBA1* WT, *UBA1* variant, pcDNA3.1 or eGPF plasmid in 20 μL Nucleocuvette Strips, using program CM137 on Amaxa 4D-Nucleofector X Unit (Lonza Bioscience). After 48 hours, cells were treated with 5 or 10 µM proteasome inhibitor MG-132; 3 or 6 µM UPR-inhibitor Ceapin-A7 (MedChemExpress); or DMSO 0.1% v/v. Cells were collected for RNA or protein isolation following 48 and 72 hours, respectively, and transfection efficiency measured by eGFP positivity using flow cytometry.

### Immunoblotting

For protein isolation, cell pellets were collected, washed in ice cold PBS and lysed in RIPA buffer (150 mM NaCl, 1% Nonidet P-40, 0.5% DOC, 0.1% SDS, 50 mM Tris pH 7.4) with cOmplete Mini Protease Inhibitor Cocktail (Roche). Proteins were quantified using Pierce BCA Protein Assay Kits (Thermo Scientific) and Infinite 200 Pro plate reader (Tecan). Samples were prepared in NuPAGE LDS Sample Buffer (4x; Invitrogen) containing 50 mM DTT, loaded on NuPAGE 4-12% Bis-Tris Gels (Invitrogen) along with PageRuler Plus Prestained Protein Ladder (Thermo Scientific) and run in NuPAGE MOPS SDS Running Buffer using XCell SureLock Mini-Cell electrophoresis system (Invitrogen). Protein was transferred on iBlot 2 Transfer Stacks (Invitrogen) and blocked in 5% non-fat dried milk in TBS-Tween. For protein detection, nitrocellulose membranes were incubated with primary antibodies overnight at 4°C followed by incubation with secondary HRP-conjugated antibodies for 1 hour at room temperature. SuperSignal West Dura Extended Duration Substrate (Thermo Scientific), Odyssey FC system and ImageStudio (Version 5.5.4, Li-Cor) were used for signal acquisition and protein quantification. Protein of interest levels were quantified relative to housekeeping protein. Antibody details are listed in Supplemental Table 2.

### Immunoprecipitation

Dynabeads Protein G Immunoprecipitation Kit (Invitrogen) was used for magnetic isolation of FLAG-tagged UBA1 from 500 µg of total protein per reaction with 5 µg of either anti-FLAG antibody or mouse IgG control. Incubation times were 30 minutes for bead conjugation and overnight at 4°C for antigen incubation. Detection of FLAG-tagged protein by immunoblotting was performed as described. Antibody details are listed in Supplemental Table 2.

### Viability and ubiquitination assays

K562 cells with a heterozygous knock-in *SF3B1*^K700E^ mutation and the parental *SF3B1*^WT^ were purchased from Horizon Discovery and cultured as above. To determine the lethal dose (LD_50_) of TAK-243 (MedChemExpress), *SF3B1*^WT^ and *SF3B1*^K700E^ cells were prepared as follows: 5×10^4^ K562 cells were seeded in 200 µL K562 medium and 4×10^4^ iPSC-HSPCs were seeded in 200 µL StemSpan SFEM II (StemCell Technologies) supplemented with 1% P/S, 50 ng/mL of IL3, SCF, FLT3L, and TPO and 10 ng/mL IL6 (PeproTech). Cells were treated with increasing doses of TAK-243 or DMSO 0.1% v/v in ultra-low attachment 96 well plates for 24 hours and assessed for viability by flow cytometry, as described below. To assess global ubiquitination upon UBA1 inhibition, *SF3B1*^WT^ K562 cells were treated with increasing doses of TAK-243 for 24 hours followed by immunoblotting for Ub-K48. ß-actin was used as loading control.

### Competitive growth co-culture of K562 cells

K562 cells were seeded at a total density of 2×10^5^ cells/mL in a 50:50 ratio of *SF3B1*^WT^:*SF3B1*^K700E^ in 24 well plates, equaling a *SF3B1*^K700E^ variant allele frequency (VAF) of 25%. Cells were treated with 50 nM TAK-243 or DMSO 0.1% v/v for 72 hours, followed by DNA extraction using QIAcube Connect with QIAmp DNA Mini Kit (Qiagen). DNA was measured by Nanodrop 2000 Spectrophotometer (Thermo Scientific) and *SF3B1*^K700E^ VAF by droplet digital PCR (ddPCR; Bio-Rad), as previously described.^32^ VAF was calculated using QuantaSoft analysis software v.1.7.4 based on Poisson distribution. Control samples with water, confirmed *SF3B1*^K700E^ and WT were included in each run.

### Colony-forming unit (CFU) assay of K562 cells

500 cells of K562 *SF3B1*^WT^ or *SF3B1*^K700E^ were seeded per 35 mm dish in Methocult H4434 (StemCell Technologies) containing 15, 32 or 50 nM TAK-243 or DMSO 0.1% v/v. Absolute colony numbers were counted after seven days using an inverted microscope.

### Flow Cytometry

Flow cytometry experiments evaluating *SF3B1*^WT^ or *SF3B1*^K700E^ iPSC-derived or K562 cells were performed on a BD LSRFortessa and analyzed with FlowJo v10.10 (BD Biosciences). Cells were stained for 30 min on ice using antibodies CD34-Pe-Cy7 and CD45-APC for hematopoietic progenitors and CD45-APC, CD71-PE and GlyA-FITC for erythroid cells with 1:200 Aqua Live/Dead viability stain (Thermo Scientific). Intracellular flow cytometry staining was prepared using Foxp3/Transcription Factor Staining Buffer Set (Invitrogen): following surface staining with CD34-PE-Cy7, cells were incubated overnight at 4°C with UBA1a antibody or matching rabbit IgG and subsequently stained with Goat anti-Rabbit AlexaFluor 488. For viability assays, cells were stained with 1:200 Aqua and 400 nM Apotracker Green (BioLegend) solution for 20 min at room temperature. Viable cells were defined as Aqua^-^/Apotracker^-^ single cells. All steps were performed in FACS Buffer (PBS + 2% FBS + 2 mM EDTA) and cells were fixed in 2% PFA prior to analysis. Antibody specifics are listed in Supplemental Table 2.

### Primary sample collection and ethical approval

Bone marrow (BM) samples were obtained from three patients with MDS-*SF3B1* and two healthy, normal BM (NBM) donors for control at Karolinska University Hospital, Huddinge, Sweden. All source material was provided with written informed consent for research use, in accordance with the Declaration of Helsinki, and the study was approved by the Ethics Research Committee at Karolinska Institutet (2017/1090-31/4, 2022-03406-02 and 2024-03119-02).

### Primary CD34^+^ CFU assay with single-colony genotyping

CD34^+^ cells were enriched from BM MNCs of sex-matched *SF3B1*-mutated MDS patients (n = 3) and healthy donors (n = 2) using CD34 Microbeads and positive selection with autoMACS Pro Separator (Miltenyi Biotec). 4000 or 7500 cells were plated as duplicates in MethoCult (H4434; StemCell Technologies) containing 32 nM TAK-243 or DMSO 0.1% v/v and cultured for 14 days. For colony counting, whole culture dishes were imaged using an Eclipse Ti2 inverted microscope (Nikon) in widefield mode and counted using cell counter plugin in Fiji (PMID: 22743772). For assessment of *SF3B1* mutation status, individual colonies were manually picked under an inverted microscope and DNA isolated using QIAamp DNA Micro Kit (Qiagen). ddPCR for *SF3B1* VAF was performed as described in main methods. Colonies with *SF3B1* VAF ≥ 40 were scored as mutant; VAF ≤ 4 as WT; and 4 < VAF < 40 as mixed and therefore excluded, see Supplemental Table 3.

### Statistical analysis

Statistical analysis was performed with Prism (v10.2, GraphPad). Data are shown as mean with standard error unless otherwise noted. Numerical variables were compared using unpaired t test (1D, 2H, 3C, 3D, 3F, 4B, 4H, S1A, S1B, S6A), one-way ANOVA (2F, 4F) and two-way ANOVA (2B, 2D, 2J, 3E, S4C). For multiple test correction, *P* values were adjusted by Šídák’s, Holm-Šídák’s, Tukey’s or Dunnett’s multiple comparisons test.

## Data availability

The raw data for the RNA sequencing and corresponding metadata and intermediate analysis has been deposited as SND-ID: 2024-128 (Bulk RNA sequencing of erythroblasts from a pair of *SF3B1*-mutated and *SF3B1*-wildtype induced pluripotent stem cell (iPSC) lines) on the Swedish National Data Service’s research data repository. These data have been uploaded with restricted access due to their personally identifiable information status but are accessible upon reasonable request through SND at: https://doi.org/10.48723/3hs1-0v44. Other raw data are available from the corresponding author upon request.

## Acknowledgments and funding

The authors thank the patients and healthy donors for their participation in this research; and the MedH Flow Cytometry Core Facility, the Live Cell Imaging Core Facility, and Science for Life Laboratory at Karolinska Institutet for experimental help and support. VL was supported by Vetenskapsrådet (grant number 2020–01902), Cancer Research KI (Karolinska Institutet), and Felix Mindus contribution to Leukemia Research; PLM was supported by Cancerfonden (grant number 21 0340) and the Myelodysplastic Syndromes foundation Inc. (grant number 1142079); EHL was supported by Cancerfonden (grant number 19 0200), Vetenskapsrådet (grant number 211133), Knut and Alice Wallenberg Foundation (grant number 2017.0359).

## Authorship

Contributions: JT, PLM and VL designed the study; JT, SH, KMK, TMB, PM conducted experiments; AGD, EP, EPP, EHL provided cells and expertise; IB and ACB assisted with biobank material; JT, SH, KMK, GT, TMB, EHL, PLM, VL analyzed and interpreted data; JT and VL wrote the manuscript, and all authors reviewed and edited the manuscript.

## Figure Legends

**Supplemental Figure 1.**
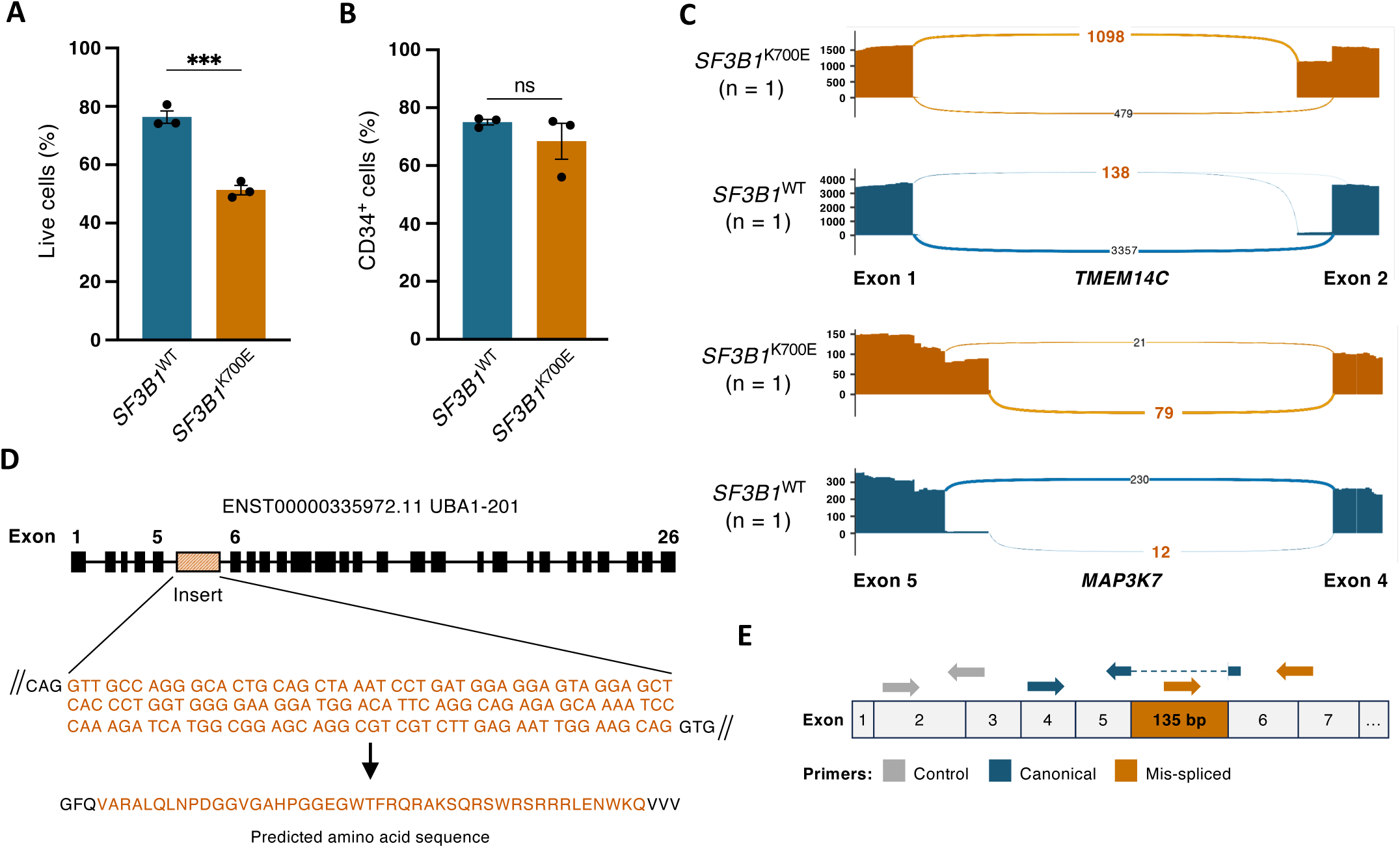
(A) Percent viable (n = 3) and (B) CD34+ cells (n = 3) from flow cytometry analyses of HSPCs from SF3B1WT and SF3B1K700E iPSCs after 12 days of differentiation, quantified from main Figure 1B. Mean ± SEM. Unpaired t test. (C) Sashimi plots of known mis-spliced regions in TMEM14C and MAP3K7 in SF3B1WT and SF3B1K700E from total RNA sequencing of iPSC-derived erythroblasts. Black, canonical splice junction counts; orange, mis-spliced junction counts. (D) Localization of UBA1 RNA mis-splicing and predicted nucleotide and amino acid sequences. (E) Primer design schematic for quantification of UBA1 splice forms by RT-qPCR. Unpaired t test; ***, P ≤ 0.001; ns, not significant.

**Supplemental Figure 2.**
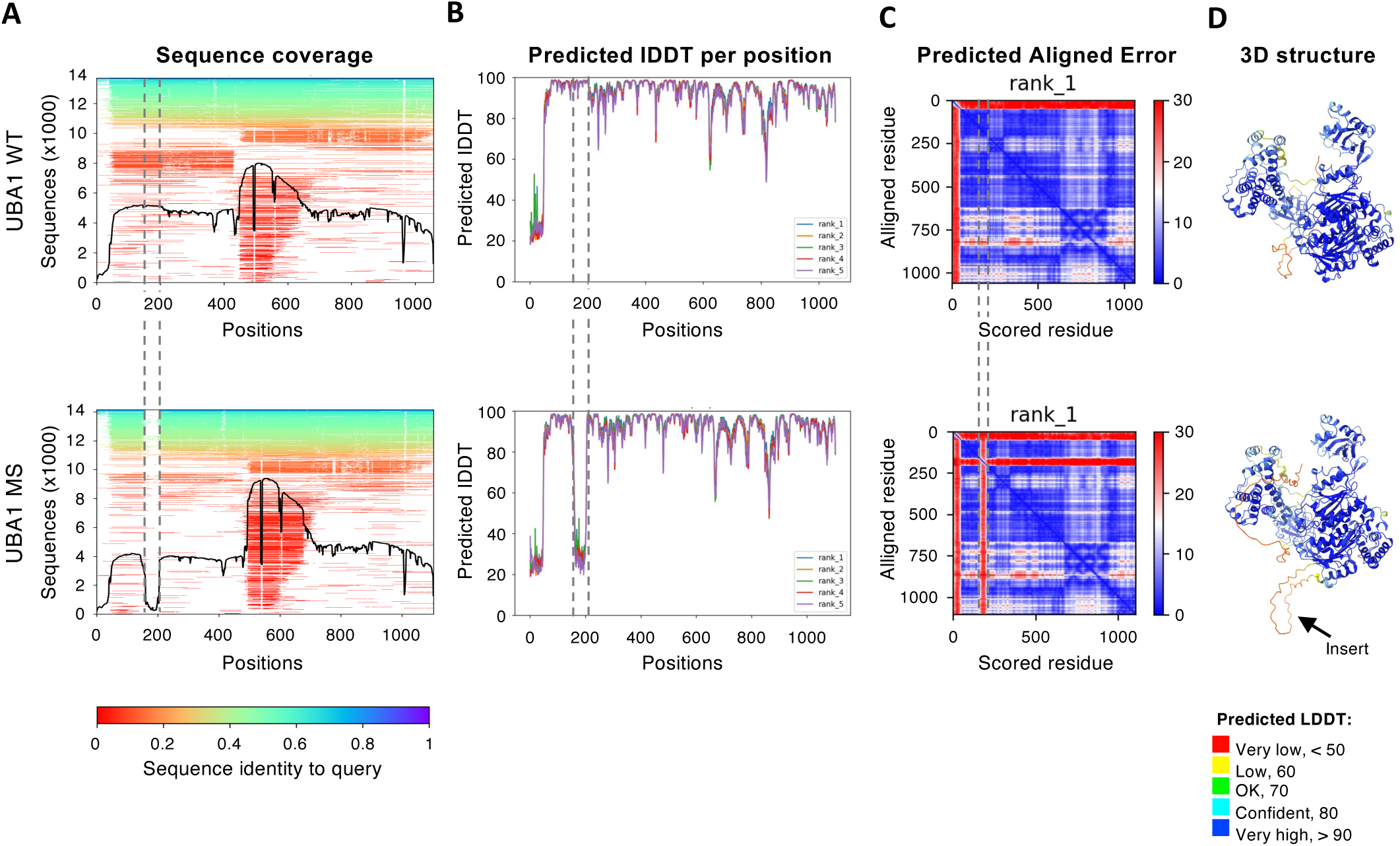
(A) Heatmap of multiple-sequence alignment (MSA) with black line qualifying relative coverage from model output of AlphaFold2 structural prediction of UBA1WT and UBA1ms amino acid sequences using ColabFold v1.5.5 (B) Predicted LDDT per residue and (C) Heatmap of predicted alignment plot for the top 5 models obtained. (D) 3D models of predicted secondary structures. Colors reflect local model confidence.

**Supplemental Figure 3.**
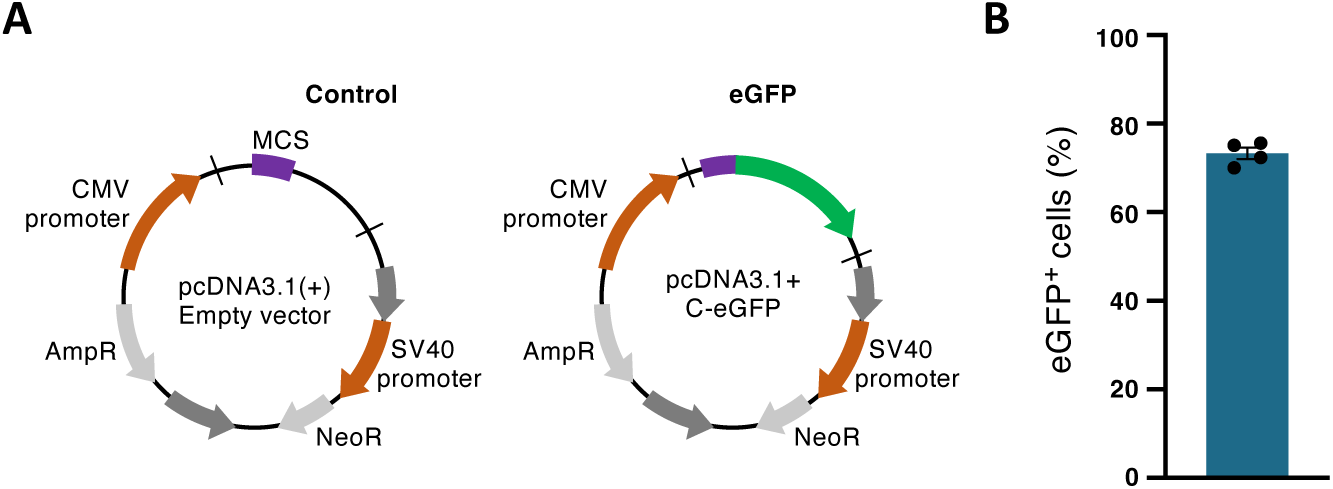
(A) Plasmid maps of empty pcDNA3.1 (control) and pcDNA3.1 C-eGFP (eGFP) vectors. (B) Percent eGFP+ K562 cells 72 hours post-transfection with eGFP control plasmid, quantified by flow cytometry (n = 4, mean ± SEM).

**Supplemental Figure 4.**
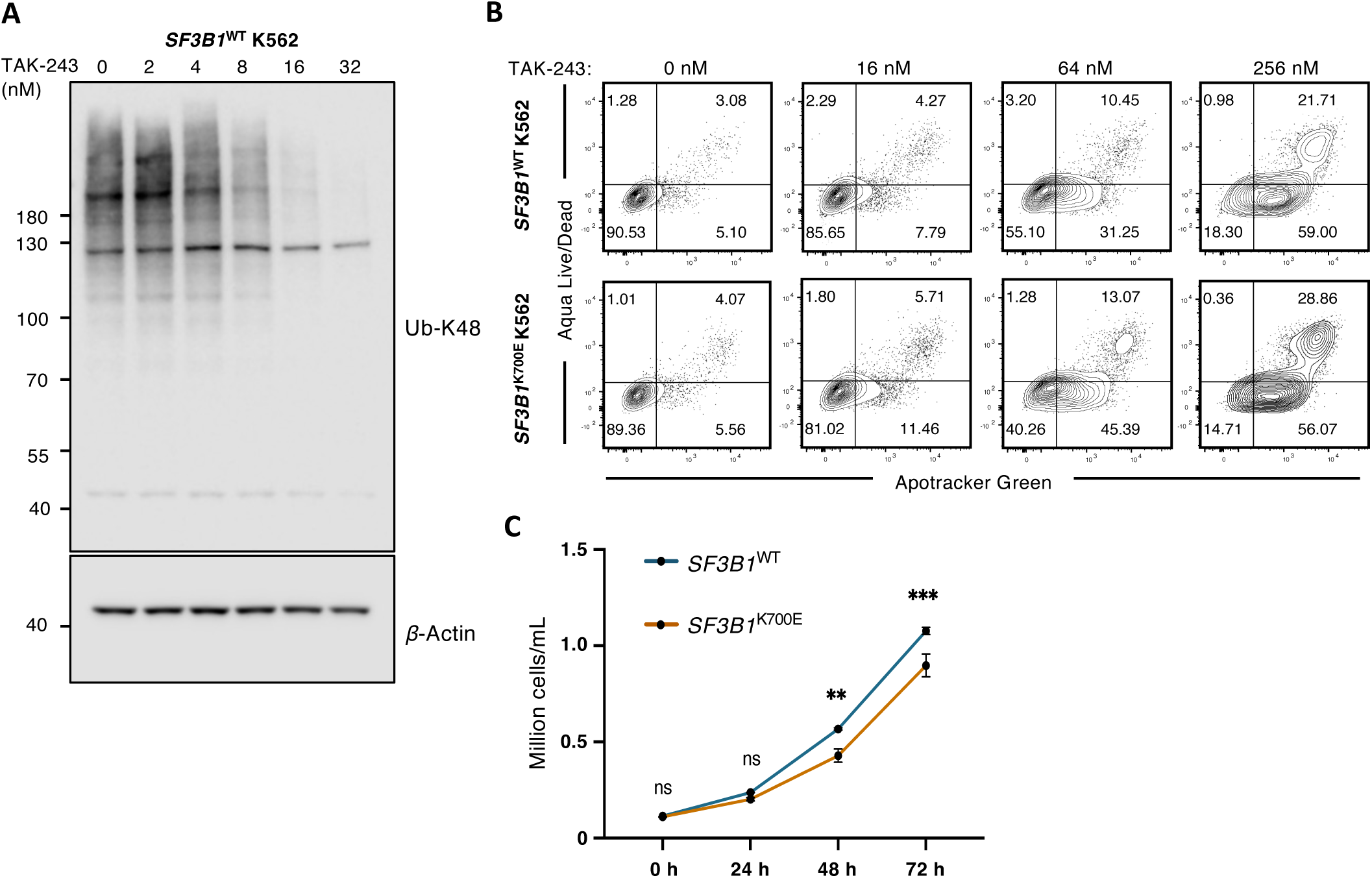
(A) Immunoblot of whole cell lysate from SF3B1WT K562 cells treated with increasing doses of TAK-243 for 24 hours. Signals for Ub-K48 and ß-actin. (B) Representative flow cytometry diagrams of SF3B1WT K562 cells treated with increasing doses of TAK-243 for 24 hours. (C) Growth curves for SF3B1WT and SF3B1K700E cells over 72 hours quantified by automatic cell counting (n = 3). Mean ± SEM. Two-way ANOVA with Šídák multiple comparisons test. **, P ≤ 0.01; ***, P ≤ 0.001; ns, not significant.

**Supplemental Figure 5.**
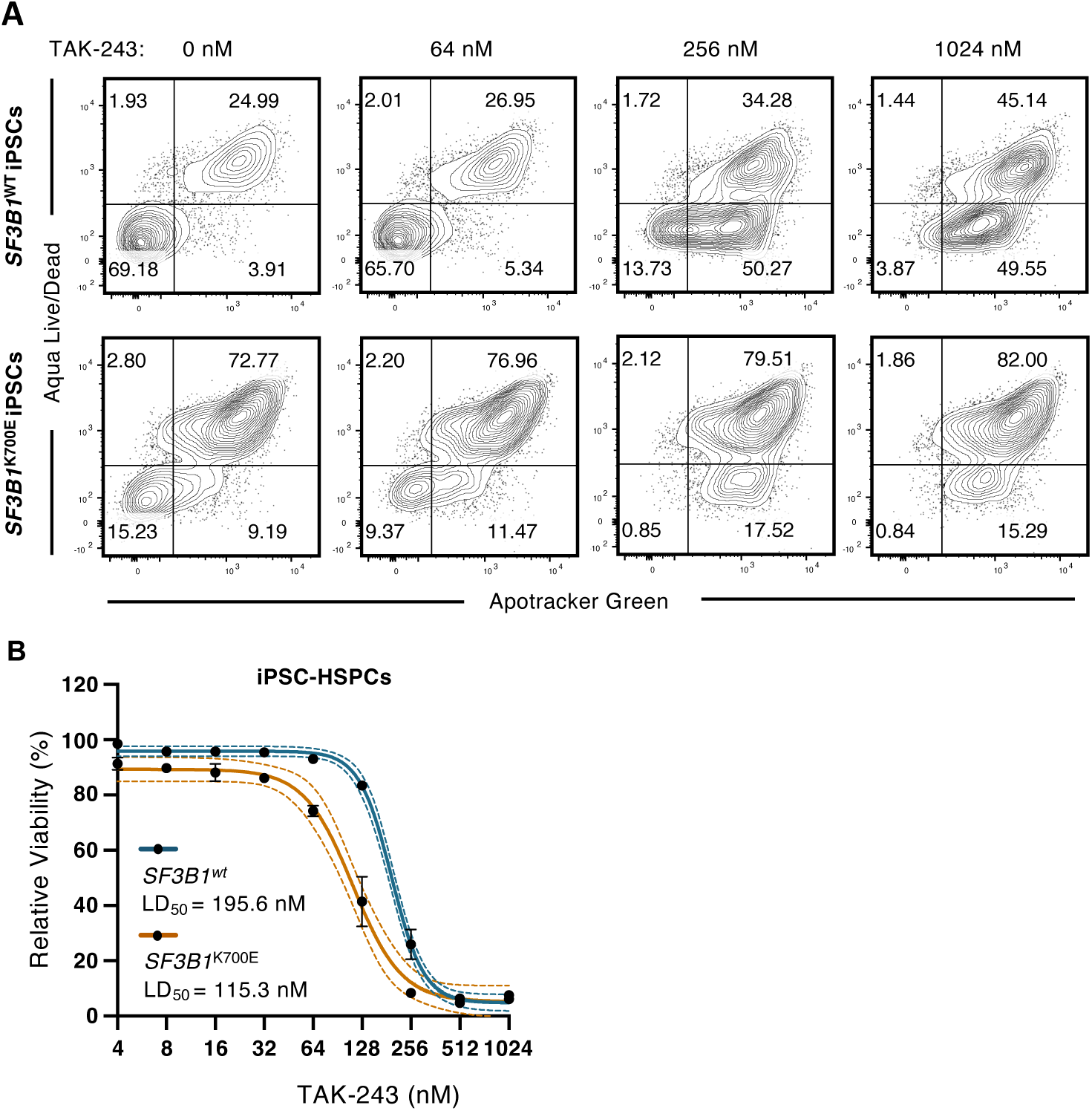
(A) Representative flow cytometry diagrams of HSPCs from SF3B1WT iPSCs treated with increasing doses of TAK-243 for 24 hours. ns, not significant. (B) Dose response curves of SF3B1WT and SF3B1K700E iPSC-derived HSPCs treated with TAK-243 or DMSO for 24 hours, analyzed as in (3B). LD50 of TAK-243 displayed under the curve are quantified in (4B) (n = 3).

**Supplemental Figure 6.**
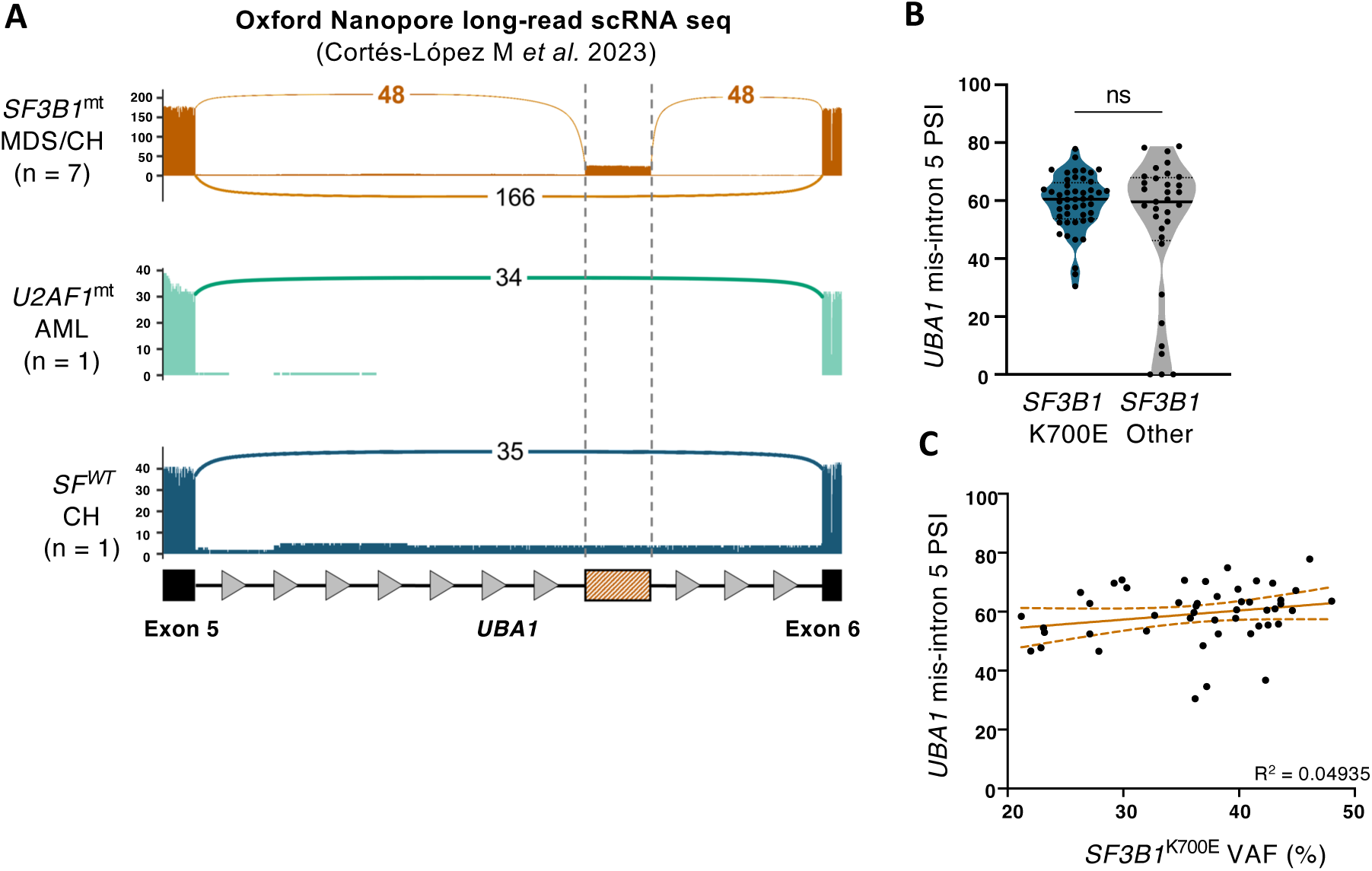
(A) Sashimi plots for read counts of the UBA1ms region from published RNA sequencing data of SF3B1mt MDS/CH (n = 7), U2AF1mt AML (n = 1) and SFWT CH (n = 1) cases.40 (B) Violin plots of PSI UBA1 intron 5 mis-splicing in SF3B1-mutated cases, comparing SF3B1K700E to all other SF3B1 variants (nK700E = 48; nOther = 33;). Unpaired t test. (C) Scatter plot of UBA1 intron 5 mis-splicing PSI over SF3B1K700E VAF in SF3B1K700E cases (n = 48). Simple linear regression (solid line) with 95 CI (dotted line). ns, not significant. CH, clonal hematopoiesis; AML, acute myeloid leukemia.

## Supplementary Tables

**Supplemental Table 1:**
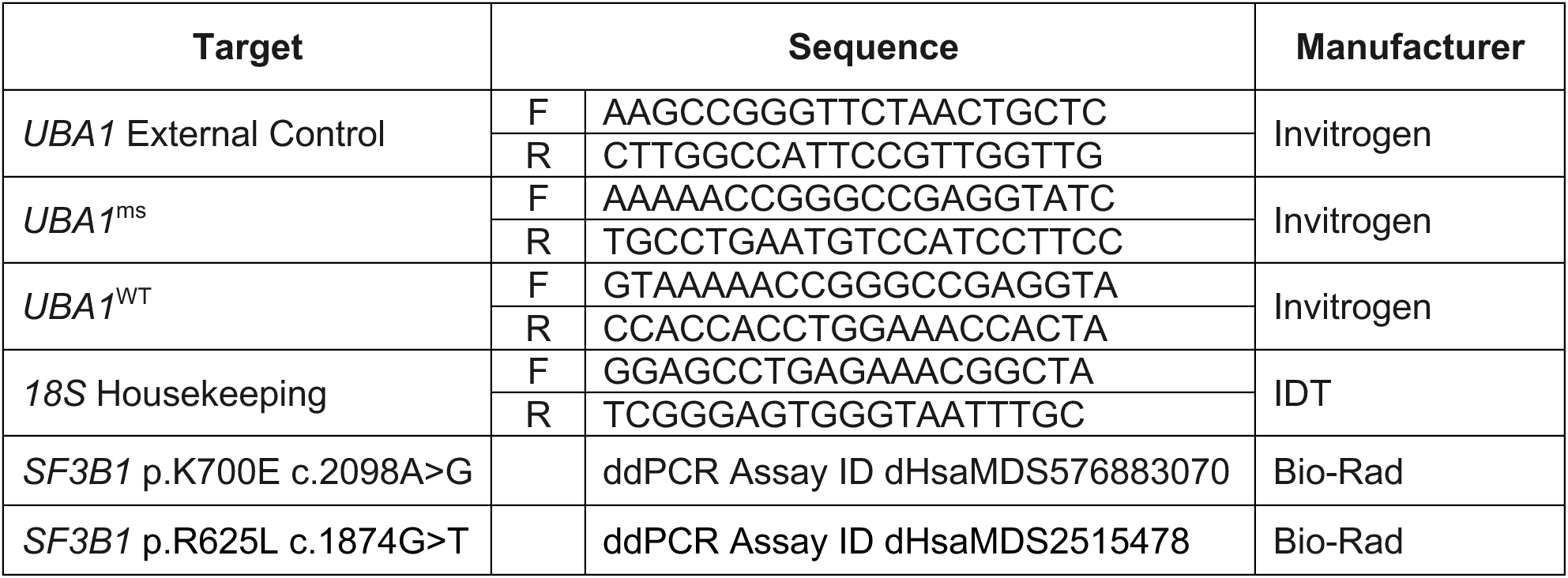
List of Primers.

**Supplemental Table 2:** List of Antibodies.

| Antigen | Conjugate | Clone | RRID | Dilution | Manufacturer |
| --- | --- | --- | --- | --- | --- |
| CD34 | PE-Cy7 | 4H11 | AB_1963576 | 1:100 | Invitrogen |
| CD45 | APC | HI30 | AB_10667894 | 1:100 | Invitrogen |
| CD71 | PE | OKT9 | AB_10717077 | 1:100 | Invitrogen |
| CD235a | FITC | HI264 | AB_10612923 | 1:50 | BioLegend |
| UBA1a | N/A | N/A | AB_10984884 | 1:50 FC; 1:1000 WB | Invitrogen |
| UBA1a/b | N/A | N/A | AB_10986095 | 1:1000 | Invitrogen |
| Ub-K48 | N/A | ARC0811 | AB_2849283 | 1:1000 | Invitrogen |
| FLAG | N/A | 5A8E5 | AB_1720813 | 1:500 | Genscript |
| Actin | N/A | C4 | AB_2335127 | 1:10000 | MPBiomedicals |
| Lamin B1 | N/A | N/A | AB_443298 | 1:1000 | Abcam |
| GaR IgG | AF 488 | N/A | AB_143165 | 1:10000 | Invitrogen |
| GaM IgG | HRP | N/A | AB_2536527 | 1:10000 | Invitrogen |
| GaR IgG | HRP | N/A | AB_2536530 | 1:10000 | Invitrogen |
| Mouse IgG | N/A | N/A | AB_2929118 | N/A | Peprtech |
| Rabbit IgG | NA | N/A | AB_2722620 | N/A | Peprtech |

**Supplemental Table 3:** Patient CFU colony genotyping.

| Patient | Treatment | CFU ID | VAF | Genotype |
| --- | --- | --- | --- | --- |
| **MDS 1**<br>*SF3B1* p.K700E<br>c.2098A>G | DMSO | 1 | 47 | mt |
|  |  | 2 | 47 | mt |
|  |  | 3 | 47.7 | mt |
|  |  | 4 | 49 | mt |
|  |  | 5 | 52.9 | mt |
|  |  | 6 | 50 | mt |
|  |  | 7 | 45.9 | mt |
|  |  | 8 | 39 | mixed |
|  |  | 9 | 49 | mt |
|  |  | 10 | 53 | mt |
|  |  | 11 | 50 | mt |
|  |  | 12 | 51.6 | mt |
|  |  | 13 | 51.4 | mt |
|  |  | 14 | 45.6 | mt |
|  |  | 15 | 48.3 | mt |
|  |  | 16 | 51 | mt |
|  |  | 17 | 46 | mt |
|  |  | 18 | 3 | WT |
|  |  | 19 | 46.9 | mt |
|  |  | 20 | 0.2 | WT |
|  |  | 21 | 45 | mt |
|  |  | 22 | 50 | mt |
|  |  | 23 | 3.9 | mt |
|  |  | 24 | 49 | mt |
|  |  | 25 | 49 | mt |
|  |  | 26 | 50 | mt |
|  |  | 27 | 40 | No amplification |
|  |  | 28 | 51 | mt |
|  |  | 29 | 58 | mt |
|  |  | 30 | 40 | mt |
|  |  | 31 | 46 | mt |
|  |  | 32 | 50.2 | mt |
|  |  | 33 | 51 | mt |
|  |  | 34 | 47.8 | mt |
|  |  | 35 | 47 | mt |
|  |  | 36 | 50.5 | mt |
|  |  | 37 | 66 | No amplification |
|  |  | 38 | 31.1 | No amplification |
|  |  | 39 | 50 | mt |
|  |  | 40 | 52.9 | mt |
|  |  | 41 | 46 | mt |
|  |  | 42 | 45 | mt |
|  |  | 43 | 49.4 | mt |
|  |  | 44 | 51.1 | mt |
|  |  | 45 | 52 | mt |
|  |  | 46 | 50 | mt |
|  |  | 47 | 44 | mt |
|  |  | 48 | 51 | mt |
|  | TAK-243 | 1 | 48 | mt |
|  |  | 2 | 39 | mixed |
|  |  | 3 | 50.3 | mt |
|  |  | 4 | 51 | mt |
|  |  | 5 | 53 | mt |
|  |  | 6 | 50.4 | mt |
|  |  | 7 | 53 | mt |
|  |  | 8 | 51 | mt |
|  |  | 9 | 21.1 | mixed |
|  |  | 10 | 1.1 | WT |
|  |  | 11 | 37.8 | mixed |
|  |  | 12 | 58 | No amplification |
|  |  | 13 | 47 | mt |
|  |  | 14 | 51 | mt |
|  |  | 15 | 48 | mt |
|  |  | 16 | 52 | mt |
|  |  | 17 | 63 | No amplification |
|  |  | 18 | 55 | mt |
|  |  | 19 | 49 | mt |
|  |  | 20 | 51 | mt |
|  |  | 21 | 49 | mt |
|  |  | 22 | 21 | mixed |
|  |  | 23 | 0.8 | WT |
|  |  | 24 | 35 | No amplification |
|  |  | 25 | 39 | mixed |
|  |  | 26 | 2.1 | WT |
|  |  | 27 | 48.1 | mt |
|  |  | 28 | 62 | mt |
|  |  | 29 | 0.13 | WT |
|  |  | 30 | 31.2 | mixed |
|  |  | 31 | 44 | mt |
|  |  | 32 | 50.3 | mt |
|  |  | 33 | 44 | mt |
|  |  | 34 | 39 | No amplification |
|  |  | 35 | 33 | No amplification |
|  |  | 36 | 47 | mt |
|  |  | 37 | 45.5 | mt |
|  |  | 38 | 51 | mt |
|  |  | 39 | 50.6 | mt |
|  |  | 40 | 53 | mt |
|  |  | 41 | 52 | mt |
|  |  | 42 | 9.8 | mixed |
|  |  | 43 | 63 | No amplification |
|  |  | 44 | 1.8 | WT |
|  |  | 45 | 49.7 | mt |
|  |  | 46 | 47 | mt |
|  |  | 47 | 3.7 | WT |
| **MDS 2**<br>*SF3B1* p.K700E<br>c.2098A>G | DMSO | 1 | 50.4 | mt |
|  |  | 2 | 49.1 | mt |
|  |  | 3 | 49.1 | mt |
|  |  | 4 | 49 | mt |
|  |  | 5 | 52 | mt |
|  |  | 6 | 51 | mt |
|  |  | 7 | 48.7 | mt |
|  |  | 8 | 48 | mt |
|  |  | 9 | 50.1 | mt |
|  |  | 10 | 52.2 | mt |
|  |  | 11 | 52 | mt |
|  |  | 12 | 49 | mt |
|  |  | 13 | 45 | mt |
|  |  | 14 | 0.02 | WT |
|  |  | 15 | 48.4 | mt |
|  |  | 16 | 51.5 | mt |
|  |  | 17 | 49.1 | mt |
|  |  | 18 | 48.2 | mt |
|  |  | 19 | 51.5 | mt |
|  |  | 20 | 48 | mt |
|  |  | 21 | 0.06 | WT |
|  |  | 22 | 0.16 | WT |
|  |  | 23 | NA | No amplification |
|  |  | 24 | 51.4 | mt |
|  | TAK-243 | 1 | 49.7 | mt |
|  |  | 2 | 48.6 | mt |
|  |  | 3 | 0.4 | WT |
|  |  | 4 | 53 | mt |
|  |  | 5 | 50.3 | mt |
|  |  | 6 | 58 | mt |
|  |  | 7 | 47 | mt |
|  |  | 8 | 41 | mt |
|  |  | 9 | 49 | mt |
|  |  | 10 | 51 | mt |
|  |  | 11 | 55 | mt |
|  |  | 12 | 0.07 | WT |
|  |  | 13 | 0 | WT |
|  |  | 14 | 51.3 | mt |
|  |  | 15 | 54 | mt |
|  |  | 16 | 45 | mt |
|  |  | 17 | 50.6 | mt |
|  |  | 18 | 53 | mt |
|  |  | 19 | 0.12 | WT |
|  |  | 20 | 51.4 | mt |
|  |  | 21 | 47.3 | mt |
|  |  | 22 | 46 | mt |
|  |  | 23 | 56 | mt |
| **MDS 3**<br>*SF3B1* p.R625L<br>c.1874G>T | DMSO | 1 | 1.2 | WT |
|  |  | 2 | 47 | mt |
|  |  | 3 | 0.5 | WT |
|  |  | 4 | 3.5 | mixed |
|  |  | 5 | 0.19 | WT |
|  |  | 6 | 47 | mt |
|  |  | 7 | 0 | WT |
|  |  | 8 | 0 | WT |
|  |  | 9 | 2.9 | WT |
|  |  | 10 | 0 | WT |
|  |  | 11 | 49.4 | mt |
|  |  | 12 | 0.5 | WT |
|  |  | 13 | 5.4 | mixed |
|  |  | 14 | 0.03 | WT |
|  |  | 15 | 1.6 | WT |
|  |  | 16 | 33 | No amplification |
|  |  | 17 | 0 | WT |
|  |  | 18 | 49 | mt |
|  |  | 19 | 60 | mt |
|  |  | 20 | 2.3 | WT |
|  | TAK-243 | 1 | 0.2 | WT |
|  |  | 2 | 0 | WT |
|  |  | 3 | 2.3 | WT |
|  |  | 4 | 8.5 | mixed |
|  |  | 5 | 9.9 | mixed |
|  |  | 6 | 10.8 | mixed |
|  |  | 7 | 37 | mixed |
|  |  | 8 | 2.8 | WT |
|  |  | 9 | 0.45 | WT |
|  |  | 10 | 0 | WT |
|  |  | 11 | 0 | WT |
|  |  | 12 | 0 | WT |
|  |  | 13 | 11 | mixed |
|  |  | 14 | 0.19 | WT |
|  |  | 15 | 0.11 | WT |
|  |  | 16 | 0.15 | WT |
|  |  | 17 | 0.019 | WT |
|  |  | 18 | 0.13 | WT |
|  |  | 19 | 3.7 | mixed |
|  |  | 20 | 2.1 | WT |
|  |  | 21 | 0.23 | WT |
|  |  | 22 | 0.6 | WT |
|  |  | 23 | 18 | mixed |

